# A microfluidic platform for characterizing the structure and rheology of biofilm streamers

**DOI:** 10.1101/2022.02.22.481486

**Authors:** Giovanni Savorana, Jonasz Słomka, Roman Stocker, Roberto Rusconi, Eleonora Secchi

## Abstract

Biofilm formation is the most successful survival strategy for bacterial communities. In the biofilm lifestyle, bacteria embed themselves in a self-secreted matrix of extracellular polymeric substances (EPS), which acts as a shield against mechanical and chemical insults. When ambient flow is present, this viscoelastic scaffold can take a streamlined shape, forming biofilm filaments suspended in flow, called streamers. Streamers significantly disrupt the fluid flow by causing rapid clogging and affect transport in aquatic environments. Despite their relevance, the structural and rheological characterization of biofilm streamers is still at an early stage. In this work, we present a microfluidic platform that allows the reproducible growth of biofilm streamers in controlled physico-chemical conditions and the characterization of their biochemical composition, morphology, and rheology *in situ*. We employed isolated micropillars as nucleation sites for the growth of single biofilm streamers under the continuous flow of a diluted bacterial suspension. By combining fluorescent staining of the EPS components and epifluorescence microscopy, we were able to characterize the biochemical composition and morphology of the streamers. Additionally, we optimized a protocol to perform hydrodynamic stress tests *in situ*, by inducing controlled variations of the fluid shear stress exerted on the streamers by the flow. Thus, the reproducibility of the formation process and the testing protocol make it possible to perform several consistent experimental replicates that provide statistically significant information. By allowing the systematic investigation of the role of biochemical composition on the structure and rheology of streamers, this platform will advance our understanding of biofilm formation.

## 1 Introduction

The bacterial colonization of surfaces is commonly associated with the release of extracellular polymeric substances (EPS), which self-assemble into a protective matrix. ^1–3^ The bacterial communities embedded in such a polymeric scaffold are called biofilms. ^4^ EPS include polysaccharides, proteins, and extracellular DNA (eDNA). ^3^ The polymer matrix provides protection against mechanical insults: its viscoelastic behavior allows for effective stress dissipation and adaptation to the persistent action of external forces. ^5–7^ The viscoelastic adaptation of biofilms often occurs in the presence of fluid flow, a ubiquitous source of mechanical stress in microbial habitats. ^8,9^ The ambient flow can shape biofilms into thin streamlined filaments, known as streamers, fixed to a tethering point, and suspended in bulk flow. ^10,11^ Streamlining allows biofilms to minimize drag, which can consequently withstand stronger flows and effectively colonize different flow environments. Streamer formation has been observed on obstacles in a flow path, such as porous media and medical devices, ^12–16^ or on objects moving in a fluid, like marine particles, rising oil droplets or sinking marine snow in the ocean. ^17,18^ Thus, streamers appear to play a crucial role both in medical and environmental settings.

Although it has already been a few decades since scientists reached a consensus on biofilms being the predominant bacterial lifestyle, their rheological investigation is still at an early stage. ^19^ The macro- and microrheological characterizations of surface-associated biofilms revealed that the parameters describing biofilm rheology, namely elastic modulus and viscosity, span orders of magnitudes in biofilms with different compositions and grown in different environments. ^6^ However, a clear link between composition, growth conditions, and the resulting biofilm properties remains to be established, especially in the case of biofilm streamers. As streamers are suspended in flow, additional experimental challenges arise, and, consequently, techniques to perform systematic and reproducible investigations are still lacking. ^10^

While traditional rheological and microrheological techniques can be applied to surface-attached biofilms, ^19^ albeit with some caveats, ^20^ this is generally not true for biofilm streamers. First, streamers must be probed *in situ*, since their formation process and integrity are controlled and maintained by the surrounding flow. Moreover, their micrometer-sized diameters make standard *in situ* microrheological techniques impractical. A promising, non-invasive technique for characterizing the rheology of streamers exploits hydrodynamic stresses exerted by fluid flow. ^21–26^ The typical experiment is carried out by growing biofilms on the walls of mesoscopic flow chambers under the continuous flow of growth media. Selected samples are then subjected to controlled perturbations of the background flow, which is used as a mechanical probe. Their time-dependent deformation is measured as a function of the applied stress, which provides a rheological characterization of biofilms. However, the estimation of the applied stress constitutes a major source of uncertainty, due to the irregular and random shape of the streamers formed in flow chambers ^21^ and their tridimensional nature. ^26^ Thus, we still lack standardized techniques to characterize the mechanical properties of streamers *in situ* and correlate these with their structures and EPS composition as well as with the growth conditions.

In this regard, microfluidic technology provides a better control on the local hydrodynamics with respect to mesoscopic flow chambers and thus improves the reproducibility of streamer formation. ^27–31^ Moreover, the precise control offered by microfluidics on the physico-chemical environment can be exploited to investigate the effect of the growth conditions on streamer formation. However, despite such advantages, microfluidic platforms have not been optimized yet to test streamer rheology.

This article presents a microfluidic platform optimized to induce the reproducible formation of biofilm streamers and to characterize their morphology, biochemical composition and rheology *in situ*. The basic unit of the device is a straight channel with an isolated pillar located at its half-width, acting as a nucleation site for streamer formation (Fig. 1). While flowing a diluted suspension of *Pseudomonas aeruginosa* PA14 bacterial cells, we observed the formation of two streamers on the sides of the pillar, with a well-defined shape and located on the mid-horizontal plane of the channel. The reproducibility of tethering points locations and the regular shape of the streamers allowed us to systematically investigate their morphology and biochemical composition via optical microscopy. Moreover, we characterized their rheology with creep-recovery tests, carried out by exposing streamers grown at a flow velocity of 2.1 mm/s to a sudden doubling of the flow velocity for 5 min. We then used 3D numerical simulations of the flow to quantify the hydrodynamic stresses exerted by the fluid flow on the streamers. Thanks to the combination of experiments and numerical simulations, we were able to quantify the viscoelastic behavior of the streamers *in situ* and with unprecedented precision. By systematically comparing different bacterial strains and growth conditions, this platform will shed a new light on what determines the structural and rheological properties of biofilm streamers.

**Fig. 1.**
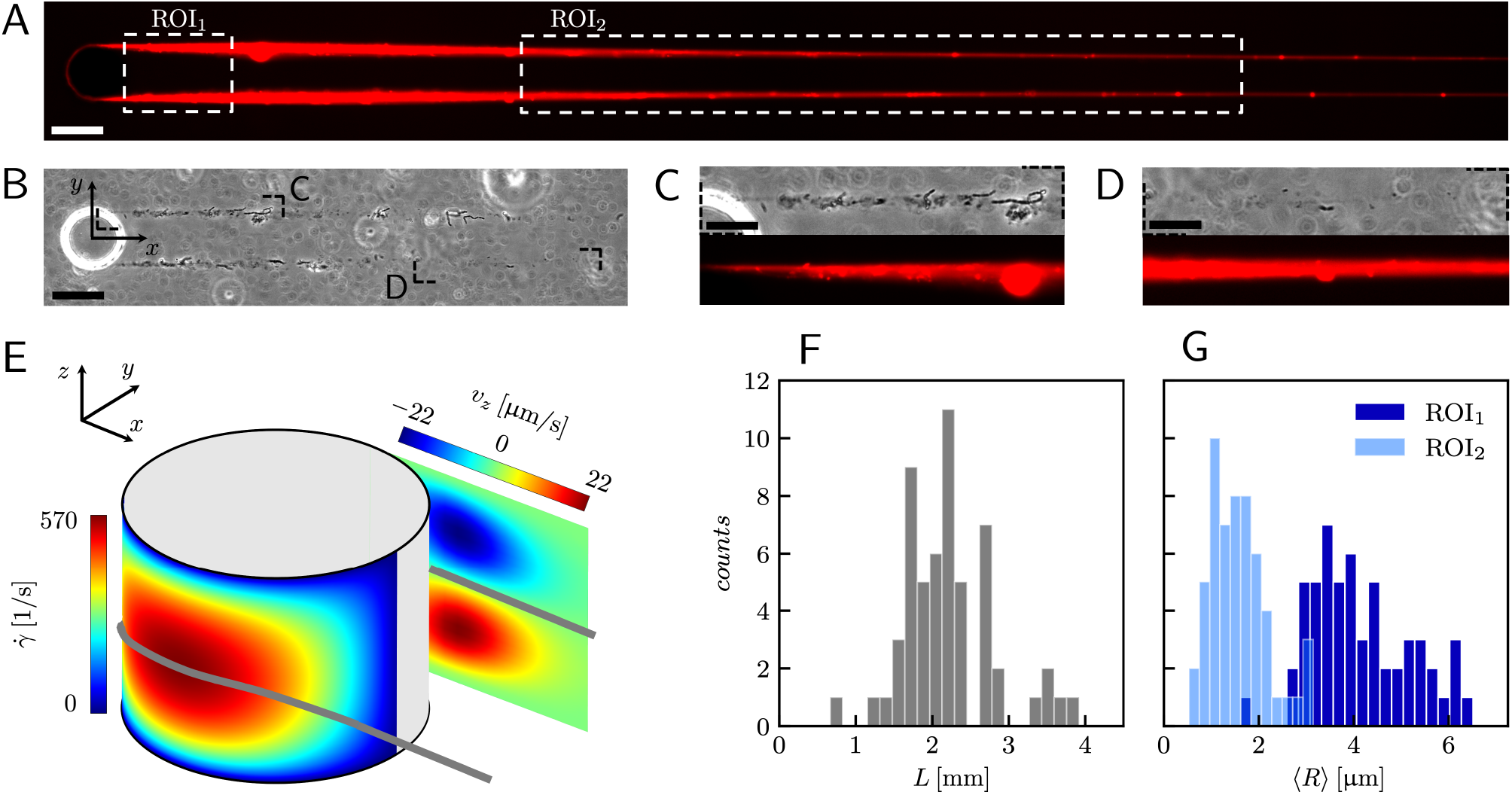
(A) Representative fluorescence and (B) phase contrast images of PA14 biofilm streamers tethered to a micropillar. The images were acquired by focusing on the channel midplane. In (A) we show the regions of interest (ROI) where we calculated the average radius 〈*R*〉 (ROI_1_ and ROI_2_) and where we acquired images during the rheological tests (ROI_2_). ROI_1_ goes 25 μm to 125 μm, while ROI_2_ goes from 400 μm to 1665.6 μm. The reported image is composed of two adjacent fields of view stitched together. ^33^ Scale bars are 50 μm. (C, D) detailed view of the two regions marked in (B). Scale bars are 20 μm. (E) Schematic of the hydrodynamic features driving streamers formation. On one side of the pillar (*y* < 0), we show the shear rate 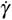 at the surface. On the other side of the pillar (*y* > 0), we show the z-component *v*_*z*_ of the velocity field on a vertical plane from *x* = 0 to *x* = 45 μm. The distributions reported here were numerically calculated in the absence of biofilm filaments and are symmetric with respect to the *x*-*z* plane. The grey lines mark the approximate position where biofilm streamers form. (F) Distributions of lengths and (G) radii (averaged over ROI_1_ and ROI_2_) of 15 h old biofilm streamers, formed in the microfluidic platform at a mean flow velocity of *U* = 2.1 mm/s. The average length is 〈L〉 = 2.22 ± 0.08 mm, while the average values of the radius are 〈R〉 = 4.1 ± 0.14 μm in ROI_1_ and 〈R〉 = 1.57 ± 0.08 μm in ROI_2_. The uncertainties on the reported values are calculated as the standard deviation of the mean.

## 2 Materials and methods

### 2.1 Microfluidic assay

To trigger the reproducible formation of biofilm streamers, we fabricated a polydimethylsiloxane (PDMS) microfluidic platform, composed of four straight channels with six isolated pillars inside each (ESI†, Fig. S1). The fabrication was carried out using standard soft lithography and PDMS molding techniques. ^32^ Each channel is 1 mm wide (W), 40 μm high (H), and 5 cm long. The cylindrical pillars have a diameter D = 50 μm. They are located at the channel half-width (*y* = 0), with a streamwise inter-pillar spacing of 5 mm. This distance ensures that the streamers tethered to a pillar do not perturb the fluid dynamic conditions of the pillar located downstream. The four channels on the platform are located 1.5 mm apart and have independent inlets and outlets. Such a parallelization allows the testing of multiple conditions during the same experimental run, which minimizes the biological variability. The flow of bacterial suspension through the channel was driven by a syringe pump (neMESYS 290N, CETONI, Germany). We used glass syringes (#81620, Model 1010 TLL, PTFE Luer Lock syringe, Hamilton Company) in order to reduce the fluidic compliance and increase the responsiveness of the system to the rapid changes in the flow rate imposed during the mechanical tests. The syringes were connected to the microchannels via dispensing needles (inner diameter 431.8μm outer diameter 635 μm, #5FVJ3, Grainger) and Tygon tubing (inner diameter 508 μm, outer diameter 1.524 mm, #AAD04103, Saint-Gobain). In this study, biofilm streamers were grown by flowing the suspensions at an average flow velocity of *U* = 2.1 mm/s for 15 h. All the experiments were performed at room temperature (T = 23±1°C). The temperature was monitored by the temperature sensor of a microscope stage top incubator (UNO-T Stage Top Incubator, Okolab).

### 2.2 Bacterial cultures

The experiments were performed using three strains of *P. aeruginosa*, a common bacterial pathogen: the PA14 wild type (WT) strain, the Pel deletion mutant PA14 Δ*pelE* and the Pel overproducer strain PA14 Δ*wspF*. All the bacterial strains were kindly provided by the laboratory of Prof. Leo Eberl at the Department of Plant and Microbial Biology, University of Zürich (Switzerland). Single colonies were grown from frozen stocks on Luria broth agar plates incubated at 37 °C for 24 h. Then, bacterial suspensions were prepared by inoculating 3 ml of tryptone broth (10 g/l tryptone, 5 g/l NaCl) with cells from a single colony and incubating at 37 °C for 3 h, while shaking at 200 rpm. The suspensions where then diluted in fresh tryptone broth to a final optical density of OD_600_ = 0.01. Biofilm streamers were visualized by fluorescently staining the eDNA (Fig. 1A) with propidium iodide (Sigma Aldrich) at a final concentration of 2 μg/ml.

### 2.3 Characterization of streamer biochemical compositions and morphology

All the images were acquired with a digital camera (ORCA-Flash 4.0 V3 Digital CMOS camera, Hamamatsu Photonics, Japan) mounted on an inverted microscope (Ti-Eclipse, Nikon, Japan) with a 20X objective magnification (CFI S Plan Fluor ELWD ADM 20XC, Nikon, Japan). Optical microscopy allowed to characterize the morphology and composition of the streamers *in situ*. Bacterial cells attached to the streamers were imaged in a phase-contrast configuration (Fig. 1B), while epifluorescence microscopy allowed the visualization of the fluorescently stained polymeric scaffold of the streamers (Fig. 1A). Since the field of view at full frame was 665.6 μm wide, several images at different downstream positions on the channel midplane were acquired to image the millimeter long streamers formed in our platform. To quantify the distribution of lengths and radii of the streamers, we acquired fluorescence images in 56 independent experimental replicates. Image analysis was performed using custom Python software. To correct for the shading artifacts resulting from the fluorescent illumination, we divided the images by a smoothed version of a calibration image acquired in a region of the platform free of any sample. The smoothing of the calibration image was performed by applying a Gaussian filter with standard deviation *σ* = 32.5 μm. We then stitched the different fields of view, ^33^ to obtain a single image of the millimeter long streamers. To perform the morphological analysis, we binarized the stitched images using a threshold intensity value 15% higher than the background intensity value. Then, we removed noise by applying an opening operation on the images, followed by a closing operation, ^34^ and we visually inspected the resulting images to eliminate the few artifacts generated by eDNA aggregates on the channel surface. Finally, we extracted the coordinates of the outline of the streamers and smoothed them with a Savitzky-Golay filter (15-μm window, 2^nd^ order polynomial). Under the assumption that the streamers lie on the channel midplane (*z* = 0) and have variable circular cross-sections with radius *R* (*x*) and center C = C(*x*, *y*_C_ (*x*), 0), we extracted the length L and the radius *R* (*x*) of each streamer from the smoothed data.

### 2.4 Creep-recovery tests on mature biofilm streamers

For the rheological characterization, creep-recovery tests were performed by imposing step-wise changes in the flow velocity of the surrounding liquid medium and by simultaneously tracking the flow-induced deformation of the streamers. The streamers were tested after 15 h of growth at a flow velocity *U* = 2.1 mm/s. We performed 10-minute-long creep-recovery tests by imposing a flow profile composed of three stages: an initial stage (0 s ≤ *t* < 150 s), a creep stage (150 s ≤ *t* < 450 s) and a recovery stage (450 s ≤ *t* < 600 s) (Fig. 2A). In the initial stage, we kept the unperturbed flow velocity constant (*U*_in_ = 2.1 mm/s); in the creep stage, we doubled the average flow velocity (*U*_cr_ = 4.2 mm/s); in the recovery stage, we lowered the velocity back to its initial value (*U*_rec_ = *U*_in_ = 2.1 mm/s). During each test, we acquired images of a portion of streamer in the region between *x* = 400 μm and *x* = 1065.6 μm (Fig. 1A, ROI_2_) at 1 fps in phase-contrast configuration. For each test, the deformation of a portion of the streamer was measured by tracking the relative displacement of two of the randomly distributed cell aggregates on the streamers (Fig. 2B, green circles). Thanks to this procedure, we identified a well-defined portion of the filament (Fig. 2B, green line) and measured its length as a function of time during the mechanical test (Fig. 2C). The aggregates were tracked using the motion tracking tools of the computer graphics software Blender. ^35^

**Fig. 2.**
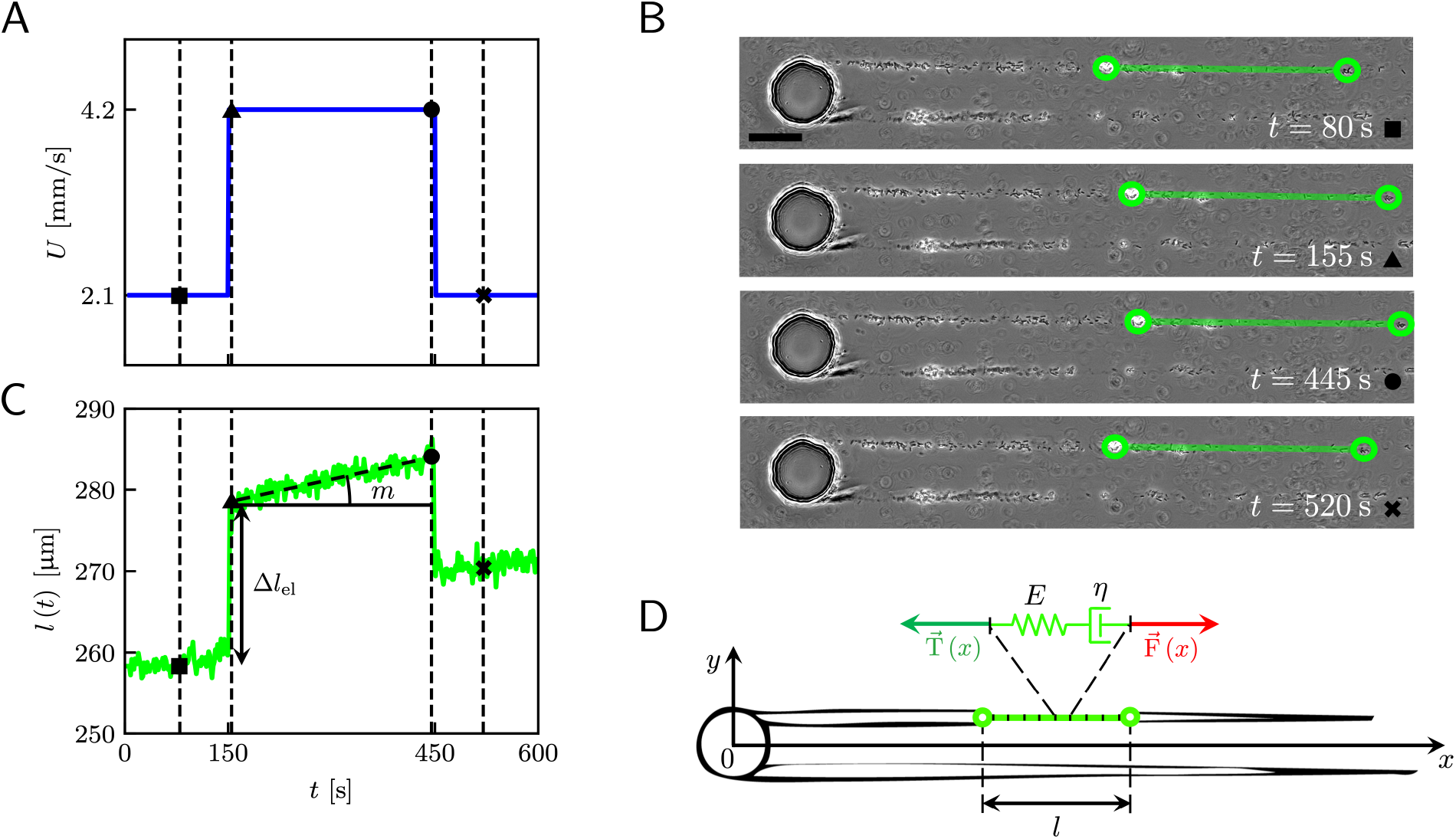
(A) Average velocity *U* in the channel as a function of time during the 10-minute creep-recovery tests. (B) Frames acquired at different stages of a mechanical test (square: 80 s; triangle: 155 s; circle: 445 s; cross: 520 s). The green circles mark the positions of two cell aggregates attached to one of the streamers, while the green line highlights the portion of streamer between them. The scale bar is 50 μm. In this experiment, the field of view was chosen to show also the pillar; the images analyzed in this work were acquired downstream, in a region corresponding to ROI_2_ of Fig. 1A. (C) Plot of the length *l* of the portion of streamer between the two aggregates in B as a function of time, which behaves as a viscoelastic fluid. Δ*l*_el_ is the instantaneous elastic deformation, while *m* is the rate of viscous deformation. The black symbols mark the values of the corresponding lengths shown in the frames of panel B. (D) The rheological behavior of any infinitesimal element of the tracked portion of the streamer can be described by a Maxwell model, with a spring with Young’s modulus *E* and a dashpot with viscosity *η* in series. The hydrodynamic force 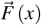 (red arrow) that stretches the infinitesimal element at x can be numerically computed. 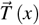 (green arrow) is the elastic reaction force, which is equal in magnitude and opposite in sign to 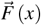.

### 2.5 Characterization of the flow field

In order to calculate the rheological properties of the streamers, we need to estimate the stresses exerted on the streamers during the creep-recovery tests. To this end, we characterized the flow field around the streamers by performing computational fluid dynamics (CFD) simulations, benchmarked with particle tracking velocimetry (PTV) experiments.

#### 2.5.1 CFD simulations

We characterized the 3D hydrodynamic conditions around the streamers for each experimental replicate by performing numerical simulations with COMSOL Multiphysics ^36^ integrated with MATLAB (LiveLink for MATLAB). MATLAB scripting with LiveLink allowed us to build 3D Comsol models of each pair of streamers, based on the morphological data obtained as explained in Section 2.3. In particular, we built a 3D loft volume for each filament, developed from cross-sections of the filaments spaced by 40 μm along the length of the streamers. As guide curves for the loft operation, we built the streamer outlines at *z* = 0: *y*(*x*) = *y*_C_(*x*) + *R*(*x*) and *y* = *y*_C_(*x*) − *R*(*x*) for each filament. The lofting was performed by interpolating the cross-sections along the guide curves, resulting in smooth 3D objects that approximate the morphological data from the experiments (ESI†, Fig. S2). The hydrodynamic problem was then solved using the Laminar Flow interface of the CFD module, with the incompressible form of the Navier-Stokes and the continuity equations. We imposed the average flow velocity value *U* at the inlet, zero outlet pressure and no-slip boundary conditions at the channel and pillar walls and on the surface of the streamers. We considered impermeable streamers. To reduce the computational time, we set the channel midplane as a symmetry plane and solved the system for the upper half of the channel. The results were then mirrored across the symmetry plane. The typical mesh consisted of approximately 3 × 10^5^ elements. Subdomains were used to build a swept mesh with finer elements in the vicinity and on the surface of the streamers (see ESI^†^). The temperature was set to T = 23 °C and water was set as the flowing fluid. The typical computational time was about 20 minutes (Processor: Intel(R) Core(TM) i7-7820X CPU @ 3.60GHz; RAM: 32GB).

#### 2.5.2 Particle tracking velocimetry

We performed PTV in selected experiments to benchmark the results of the numerical simulations. In the PTV experiments, we first flowed the bacterial suspension for 15 h; once streamers were formed, we started flowing a suspension of polystyrene tracers of diameter *d*_tr_ = 1 μm (PS-Research Particles, Microparticles GmbH), at a volume fraction *ϕ* = 0.25 %. The rapid switch between the two flowing suspensions was performed by using a Y connector (P-514, IDEX) located before the channel inlet. This procedure allowed us to avoid contact between the cells and the tracers during the formation of the streamers. To characterize the flow field around the entire streamers, we acquired time-lapse videos of 1300 frames on the channel midplane, at different locations along the flow direction. We used bright field microscopy with the condenser diaphragm completely open (NA = 0.52), in order to minimize the depth of field. The frame rate of the camera was 800 fps and the frame size was 2048 px × 256 px. Before applying the PTV algorithm to the acquisitions, we preprocessed the images to subtract the static background, calculated by averaging the intensity of the whole image stack. We segmented the particles in each frame by applying an intensity threshold equal to half the minimum intensity value calculated in each frame. Particle tracking was performed on the segmented images with a custom software based on the Trackpy Python package. ^37^ Thanks to the small depth of field, combined with image segmentation and filters on feature size, we selected particles lying on the midplane with a spread in the *z* direction of about 2*d*_tr_.

## 3 Results

### 3.1 Biochemical composition and morphology of the streamers

A continuous flow (*U* = 2.1 mm/s, Re = *ρU* D ≈ 0.1) of a diluted suspension of *P. aeruginosa* PA14 WT around an isolated pillar triggers the reproducible formation of a pair of streamers (Figs. 1A and 1B). The two streamers have distinct tethering points, respectively located on the side surface of pillar at (0, −D/2, 0) and (0, D/2, 0) (ESI†, Fig. S1). The streamers grow longer and thicker with time, until approximately 15 h. Then, they reach a stable configuration, where no major structural changes are observed. To a good approximation, the length *L* and the radius *R* (*x*) of mature streamers are constant within tenths of minutes (ESI†, Fig. S4), the timescale of the structural and rheological characterization procedure presented in this paper. This allows us to neglect deformations of the streamers under the action of the base flow at an average flow velocity *U* = 2.1 mm/s during the experiments. Previous works showed that streamer formation is driven by the interplay between the hydrodynamic features of the microenvironment and the rheological and self-assembly properties of the EPS. ^10^ CFD simulations of flow in our platform confirmed that the typical hydrodynamic features promoting streamer formation are present in our geometry (Fig. 1C). The first feature is a secondary flow in the *z* direction: simulations show vortices in the proximity of the tethering points, which point towards the midplane (*z* = 0). Such a secondary flow promotes the accumulation of EPS and cells at half-height of the pillar. ^28,29^ The second feature is the high flow shear nearby the surface of the pillar, ^38^ which extrudes the cell and EPS aggregates attached to the pillar. Additionally, we point out that the interplay between local flow and bacterial motility further enhances the colonization of the pillar. ^39^ Once the initial structure is formed, cells and EPS suspended in the bulk flow are captured by the streamers, ^40^ and contribute to their growth. Mature streamers are millimeter-long filaments, able to withstand the hydrodynamic stresses exerted by the ambient flow. The polymeric scaffold of the streamers was visualized with epifluorescence microscopy by using propidium iodide to stain the eDNA, which is a main component of PA14 streamers ^16^ and can be used for determining their morphology (Fig. 1A; ESI†, Fig. S6). Phase-contrast imaging showed that bacterial cells were found up to about *x* = 1 mm from the pillar and were more numerous in the first few hundreds of micrometers (Figs. 1C and 1D). The comparison between phase-contrast and fluorescence data shows that the EPS scaffold of the streamers had a non-negligible radius even where no cells were present. By applying the morphological analysis described in Section 2.3 to 56 independent experimental replicates, we characterized the distribution of lengths (Fig. 1F) and radii (Fig. 1G) of PA14 WT streamers after 15 h of continuous flow. The average length of the filaments was 〈*L*〉 = 2.22 ± 0.08 mm. The radius decreased with *x*: the average value in the region between *x* = 25 μm and *x* = 125 μm (Fig. 1A, ROI_1_) was 〈*R*〉 = 4.1 ± 0.14 μm and in the region between *x* = 400 μm and *x* = 1665.6 μm (Fig. 1A, ROI_2_) was 〈*R*〉 = 1.57 ± 0.08 μm. The uncertainties on the reported values are calculated as the standard deviation of the mean.

### 3.2 Creep-recovery tests

When subjected to a stress increase, streamers behave as viscoelastic fluids, undergoing an instantaneous elastic deformation, described by the Young’s modulus *E*, followed by a viscous flow, characterized by an effective viscosity *η* (Fig. 2). Our platform allows the measurement of *E* and *η* of a given portion of streamer from the deformation curve acquired during the creep-recovery test. The tests were performed by imposing a controlled perturbation to the average flow velocity *U* (Fig. 2A). By tracking two cell aggregates attached to a streamer (Fig. 2B, green circles), we identify a well-defined portion of filament (Fig. 2B, green lines) and measure its length *l* (*t*) as a function of time (Fig. 2C). The deformation at the unperturbed flow velocity *U*_in_ during the initial stage (0 *s* ≤ *t* < 150 *s*) is negligible. When doubling the flow velocity to *U*_cr_ = 2*U*_in_ during the creep stage of the test, the streamers undergo an instantaneous elastic deformation 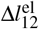 and a viscous deformation 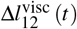, linearly increasing with time at a rate *m* (Fig. 2C). When the initial flow velocity is restored to *U*_in_ during the recovery stage, the elastic contribution to the deformation is recovered, while the viscous one is retained, due to its irreversible, dissipative nature.

Experiments show that the elastic contribution to the strain is instantaneous, whereas the viscous one is related to the progressive elongation in the time interval 150 *s* ≤ *t* < 450 *s* (Fig. 2C). Thus, we describe the behavior of any infinitesimal element of filament during our tests by using a linear viscoelastic Maxwell model, i.e. a spring and a dashpot connected in series (Fig. 2D). By adopting this model, we use the linear viscoelasticity theory, based on the assumption of small deformations. The elastic behavior of the biofilm matrix is thus described by the Young’s modulus *E* of the spring, while the viscous behavior by the viscosity *η* of the dashpot. According to the Maxwell model, after a time *t* from an axial stress increase Δ*σ* (*x*), the strain of the infinitesimal element of streamer at *x* can be written as:

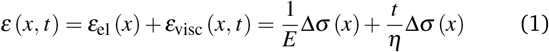

where *ε*_el_(*x*) is the elastic contribution to the strain and *ε*_visc_(*x*, *t*) is the viscous one. In this expression, we assumed that the element in the initial stage is at equilibrium, and that its viscous deformation is determined only by the stress increase Δ*σ* (*x*). This is equivalent to the assumption that the streamer behaves as a yield stress fluid, which starts flowing once the axial stress is higher than a yield stress value equal to *σ*_in_ (*x*). By integrating Eq. (1), we can write the deformation Δ*l*_12_(*t*) of the portion of streamer between two arbitrary positions *x*_1_ and *x*_2_ as:

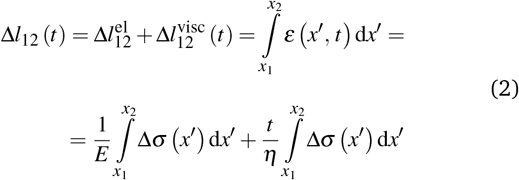

Equation (2) is equivalent to summing up the deformations of all the infinitesimal Maxwell elements composing the portion of streamer between *x*_1_ and *x*_2_. Here we assumed that all the infinitesimal Maxwell elements connected in series between *x*_1_ and *x*_2_ have the same *E* and *η*, but can be locally subjected to different stresses 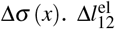 is the total elastic deformation, while 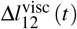 is the viscous deformation at time *t*, which can be measured from the experimental deformation curves acquired during the creep-recovery tests (Fig. 2C). Thus, *E* and *η* can be expressed as a function of the axial stress increase Δ*σ* (*x*) and the measured strain Δ*l*_12_ (*t*). In particular, the Young’s modulus of the portion of streamer between *x*_1_ and *x*_2_ is calculated as

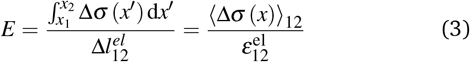

and its effective viscosity as

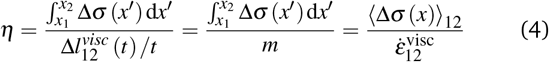

These equations quantify the average rheological properties of a portion of a streamer in terms of a series of simple one-dimensional Maxwell elements. The denominators in Eqs. (3) and (4) represent the elastic strain 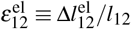 and the viscous strain rate 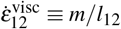 along the *x* direction, respectively, of the portion of streamer between *x*_1_ and *x*_2_. In both Eqs. (3) and (4), the numerator is the hydrodynamic axial stress increase during the creep stage of the test, averaged on the portion of the streamer between *x*_1_ and *x*_2_. This quantity has to be carefully calculated in order to obtain a reliable characterization of the streamer rheology. Thus, precise estimates of the axial force 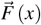 (Fig. 2D, red arrow) and of the corresponding axial stress *σ* (*x*) exerted along the portion of the streamer are needed, both during the initial and the creep stages. In section 3.3, we will present how to calculate the axial force *F* (*x*) and the corresponding axial stress *σ* (*x*) at each position *x* along the streamer. A precise estimate of the hydrodynamic stresses can be obtained with CFD simulations of flow past the streamers (Fig. 3B). The 3D models used in the simulations are built from the fluorescence images of the streamers, according to section 2.5.1. In Eqs. (1–4) we made the assumption that the axial stress step Δ*σ* (*x*) applied during the creep stage of the test is not time-dependent. In addition, we used the initial positions *x*_1_ and *x*_2_ as bounds of integration in Eqs. (2–4). As discussed in the following, these approximations are valid as long as fluid-structure interaction (FSI) is negligible, namely for small deformations of the streamers. In section 3.4, we use these approximate equations to calculate the rheological properties of *P. aeruginosa* WT streamers, without taking FSI into account. However, during the creep stage the axial stress *σ* (*x*) does not change only because the average flow velocity *U* increases, but also because the filaments are stretched and the surface exposed to flow is deformed. In section 3.5, we will take this effect into account and provide a method to estimate the impact of FSI on the axial stress *σ* (*x*) during the creep stage of the test.

**Fig. 3.**
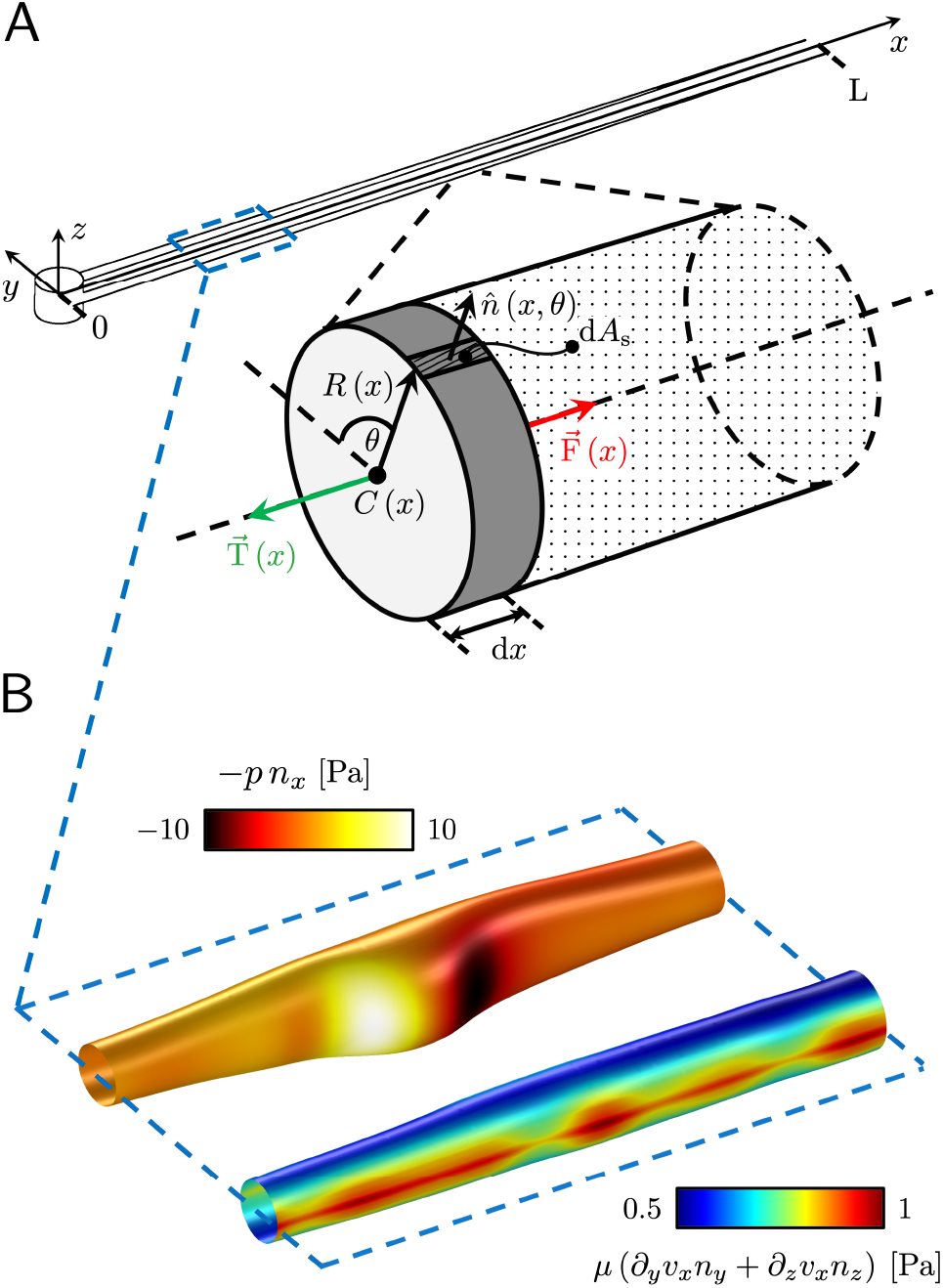
(A) Force diagram for an arbitrary infinitesimal element of biofilm streamer of cylindrical shape (shaded cylinder) located at *x*, with thickness d*x* and radius *R* (*x*). Each point on the lateral surface of the element is determined by the coordinates (*x*, *θ*) according to Eq. (5). d*A*_S_ is the infinitesimal surface element at (*θ*, *x*) and 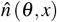 is the normal to the surface at (*θ*, *x*). Besides being subjected to local hydrodynamic stresses exerted on its lateral surface, the infinitesimal disk element has to bear the load of the whole portion downstream of *x* (dotted region, extending until *x* = *L*). 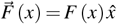 is the total axial force acting in the downstream direction (red arrow). 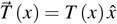 (green arrow) is the elastic reaction force, which is equal in magnitude and opposite in sign to 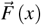. (B) Details from the results of the CFD simulation for the filaments shown in Fig. 1A. The distributions of *pn*_*x*_ (thermal colormap) and of *μ* (*∂_y_v_x_n_y_* + *∂_z_v_x_n_z_*) (rainbow colormap) are plotted on the upper filament (*y* > 0), and on the lower filament (*y* < 0) respectively, in the region between *x* = 90 μm and *x* = 190 μm.

### 3.3 Hydrodynamic forces on the streamers at equilibrium

Mature biofilm streamers are constantly subjected to a hydrodynamic force that pulls them downstream and keeps them suspended in the bulk of the flow. As pointed out in section 3.2, the streamers can withstand the force exerted by the base flow velocity *U* = 2.1 mm/s with negligible deformation. In this section, we provide an expression for the hydrodynamic force 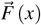 at equilibrium and for the corresponding axial stress *σ* (*x*) as a function of the hydrodynamic stresses locally exerted at the surface of the streamers. The hydrodynamic stresses are then calculated with CFD simulations, based on detailed 3D models of the streamers, according to section 2.5.1 (Fig. 3B). By comparing the numerical results and the experimental flow field obtained with PTV, we benchmark the simulation procedure (Fig. 4). The expression for the axial stress at the base flow rate *σ*_in_ (*x*) will be the starting point to describe the axial stress *σ*_cr_ (*x*) exerted during the creep stage of the test, both in the approximation of negligible FSI (section 3.4) and in the coupled case (section 3.5), where we estimate the effect of FSI. The estimates of the stress difference Δ*σ* (*x*) = *σ*_*cr*_ (*x*) − *σ*_*in*_ (*x*) between the initial and creep stages will then be used to calculate *E* and *η* from creep-recovery tests.

**Fig. 4.**
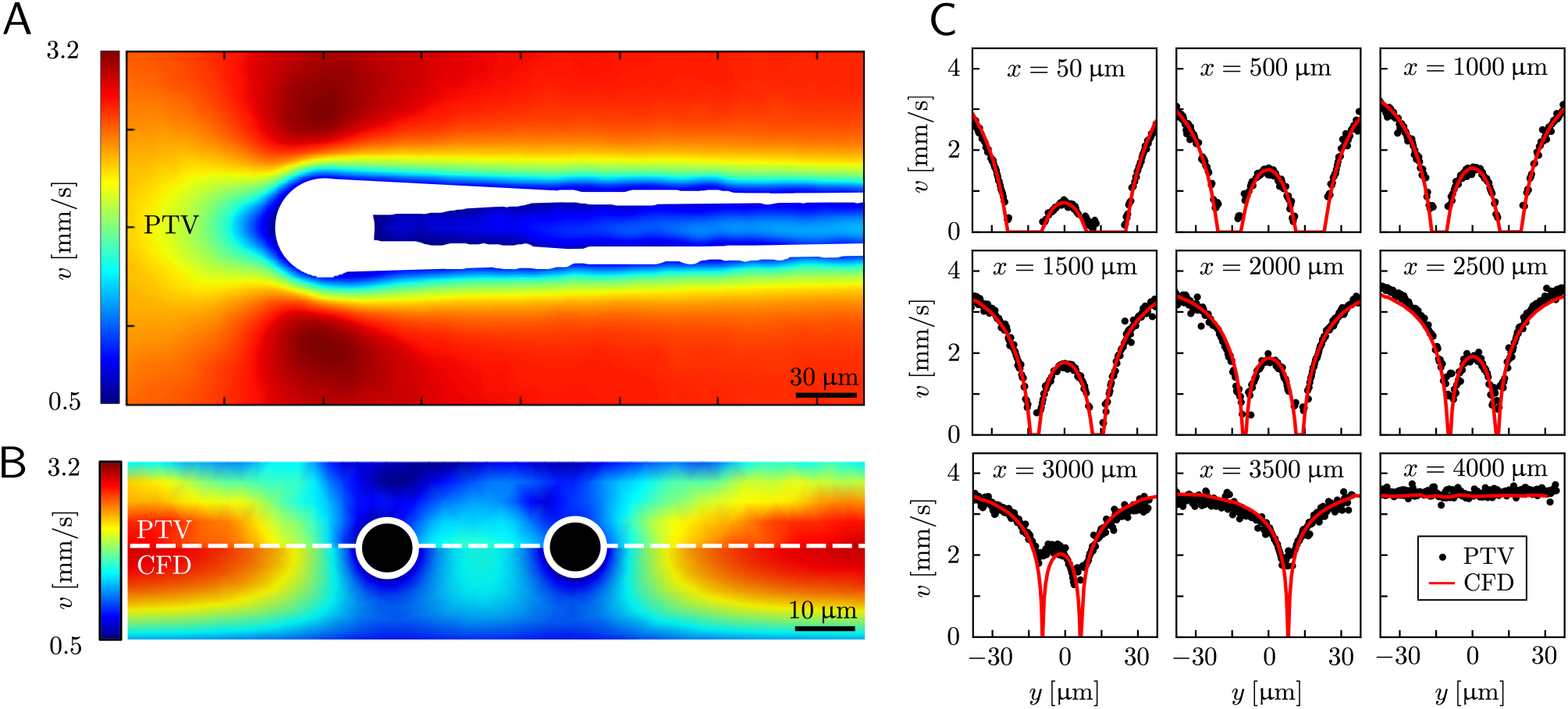
(A) Experimental map of the velocity magnitude on the channel midplane obtained with PTV. The streamers add no-slip boundaries in the middle of the channel, which drastically disrupt the flow in the channel. The map was obtained by interpolating unstructured PTV data on a regular grid of square elements with 1-μm side and by applying a gaussian smoothing with *σ* = 4 μm. (B) Experimental (upper half) and numerical (lower half) maps of the velocity magnitude on a vertical plane perpendicular to the direction of flow at *x* = 400 μm. The black circles mark the position of the cross sections of the streamers. The map was obtained by stacking 12 acquisitions taken at different heights inside the channel (Δ*z* = 2.66 μm). The 3D velocity field was averaged in the *x* direction on a 70 -μm wide region, providing a 2D velocity map on the *y*-*z* plane. The 2D velocity map was then interpolated on a regular grid of square elements with 1-μm side and smoothed with a gaussian filter with *σ* = 1 μm. (C) Experimental (black circles) and numerical (red lines) velocity profiles on the channel midplane as a function of the spanwise coordinate *y* at different downstream positions *x*. At *x* = 3500 μm only the longest filament is present, while at *x* = 4000 μm no filament is perturbing the flow field. The experimental velocity profiles are raw PTV data averaged on bins with width 1.95 μm in the *y* direction and 32.5 μm in the *x* direction, with no further processing.

We consider two mature biofilm streamers tethered to a micropillar as elastic filaments at mechanical equilibrium, subjected to hydrodynamic forces exerted by the surrounding flow. We locate the origin of the frame of reference on the vertical axis of the pillar, at half-height of the channel (ESI†, Fig. S1). We assume that each streamer is suspended on the midplane of the channel (*z* = 0), approximately parallel to the x axis, with length *L* and variable circular cross section with radius *R*(*x*), centered at C(*x*) = (*x*,*y*_C_ (*x*),0) (Fig. 3A). Thus, the shape S of each filament is represented by the following parametrization:

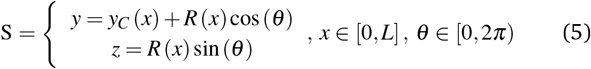

In general, the length *L* and the specific form of *R* (*x*) and C(*x*) can be different for the two streamers. The tethering points of the two streamers on the pillar are located at (0, D/2, 0) and (0, −D/2, 0) respectively. Since the streamers are approximately parallel to the *x* axis, we have that *y*_*C*_ (*x*) varies slowly with *x* (d*y*_*C*_ (*x*)/d*x* ≪ D/*L*), and further *y*_*C*_ (*x*) > 0 for one of the two filaments and *y*_*C*_ (*x*) < 0 for the other, for all *x*. Given that the streamers are aligned with the flow direction, we assume that the *x* component of the force is the one determining the axial stress that pulls the filaments downstream. We thus assume 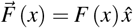. Under the assumption of mechanical equilibrium, the *x* component *T* (*x*) of the elastic reaction force exerted by the portion of the streamer upstream of *x* is *T* (*x*) = −*F* (*x*). In order to find an expression for *F* (*x*), we consider an infinitesimal disk element located at *x* with thickness d*x* and radius *R* (*x*) (Fig. 3A, grey disk in the inset). At equilibrium, the resultant force exerted on the disk is represented by the following integral from *x* to *L*:

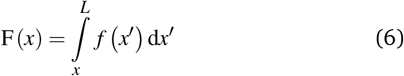

where *f* (*x*′) d*x*′ is the infinitesimal hydrodynamic force locally exerted by the fluid on the lateral surface of the infinitesimal element at *x*′. Under the approximation of weak dependence of *R* (*x*) on *x*, *f* (*x*) d*x* can be written as:

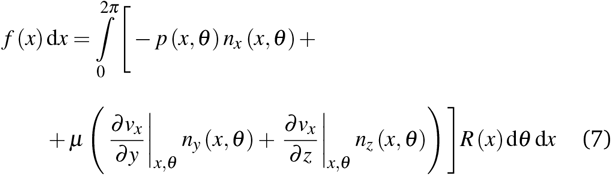

Here *n*_*i*_ (*θ*, *x*) is the *i*-th component of the normal 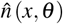 to the disk surface at 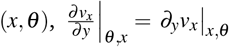 and 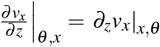 are the *xy* and *xz* components of the velocity gradient 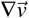 respectively, evaluated at the surface of the filament, and *μ* is the dynamic viscosity of the liquid flowing around the filaments. The expression of the force in the case of arbitrary dependence of *R* (*x*) on *x* is reported in the section 1 of the ESI†. According to Eq. (6), the axial stress on the cross section of the disk at *x* is:

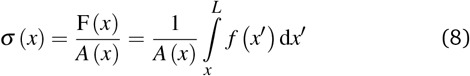

where *A* (*x*) = *πR*^2^ (*x*) is the cross sectional area of the streamer at *x*. In order to calculate the axial stress acting on the infinitesimal element at *x* according to Eq. (8), we need to quantify *pn*_*x*_, the stress contribution related to pressure, and *μ*(*∂*_*y*_*v*_*x*_*n*_*y*_ + *∂*_*z*_*v*_*x*_*n*_*z*_), the one related to shear stresses at the surface of the streamer, from *x* to *L*. Their distributions at a given flow rate depend on the detailed morphology of the filaments. As a consequence, they have non-trivial dependencies on the position. Thus, to quantify the values of *pn*_*x*_ and *μ*(*∂*_*y*_*v*_*x*_*n*_*y*_ + *∂*_*z*_*v*_*x*_*n*_*z*_) at the surface of the streamers for each experimental replicate, we performed CFD simulations of the flow based on the morphological data obtained from the fluorescence signal of the eDNA scaffold (Fig. 3B), as explained in Section 2.5.1. We point out that *pn*_*x*_ is relevant in regions where *n*_*x*_ is not significantly smaller than 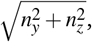, that is in regions where there is a non-negligible variation of *R* with *x*, which induces the flow lines to bend. In Fig. 3B, for example, we can see a maximum and minimum of *pn*_*x*_, upstream and downstream of a local maximum of *R* (*x*), respectively. On the other hand, *μ*(*∂*_*y*_*v*_*x*_*n*_*y*_ + *∂*_*z*_*v*_*x*_*n*_*z*_) is usually non-negligible regardless of the behavior of *R* (*x*). This term has typically a maximum on the midplane, which is the farthest away from the no-slip boundaries at the top and bottom walls of the channel. Typically, the streamers have an irregular shape with significant variations of *R* (*x*) only in the first 400 μm downstream of the pillar (Fig. 1A, ROI_1_). Elsewhere, *R* (*x*) changes slowly with *x* and 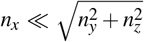. Consequently, the main contribution to the axial stress in the region investigated during the creep-recovery tests (Fig. 1A, ROI_2_) is the one related to shear stresses, while the contribution of pressure is typically 3 orders of magnitude smaller. The PTV experiments performed benchmark the numerically computed flow fields (Fig. 4). Using the experimental configuration and the analysis procedure presented in section 2.5, velocities could accurately be measured up to about 1.5 μm from the surface of the streamers. PTV results on the *x*-*y* midplane (Figs. 4A and 4C) and on the *x*-*z* vertical plane at *x* = 400 μm (Fig. 4B) are in agreement with the data obtained with the numerical simulations. This observation confirms that, within our resolution, the flow field is compatible with no-slip boundary conditions and the assumption of impermeable streamers. The streamers have a strong impact on the hydrodynamic conditions in the channel, by adding no-slip boundaries in the bulk of the flow (Fig. 4). According to CFD numerical results, the drag force exerted by the flow in the *x* direction on the biofilm-free pillar is *F*_d,*x*_ = −5.6 nN. In the case of the streamers shown in Fig. 4C, the drag force is *F*_d,*x*_ = −40.1 nN, about one higher.

### 3.4 Rheological characterization without fluid-structure interaction

As a first approach to the calculation of the axial stress *σ*_cr_ (*x*) during the creep stage, we neglect the effects of the deformation of the streamers on the flow field, so we do not take into account FSI. We use the initial undeformed geometries 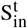, 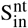 of the tracked (t, upper filament in the example reported in Fig. 2B) and non-tracked filaments (nt, lower filament in Fig. 2B) respectively in order to calculate the axial stress in both the initial (Fig. 5A) and creep stages (Fig. 5B), according to Eq. (8). During the initial stage, the axial stress as a function of *x* is:

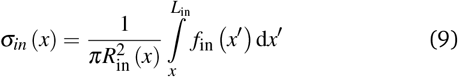

where *f*_in_ (*x*) is calculated by evaluating Eq. (7) in the initial configuration. Since we neglect FSI and the flow is in the low Re regime, the relation between the hydrodynamic stresses in the initial and creep stages is the same as the one between the average velocities. In particular, since *U*_cr_ = 2*U*_in_, we can write *f*_cr_ (*x*) = 2 *f*_in_ (*x*), and the axial stress during the creep stage can be written as:

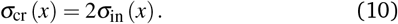

Thus, the stress step applied during the creep part of the test is:

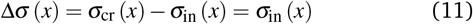

**Fig. 5.**
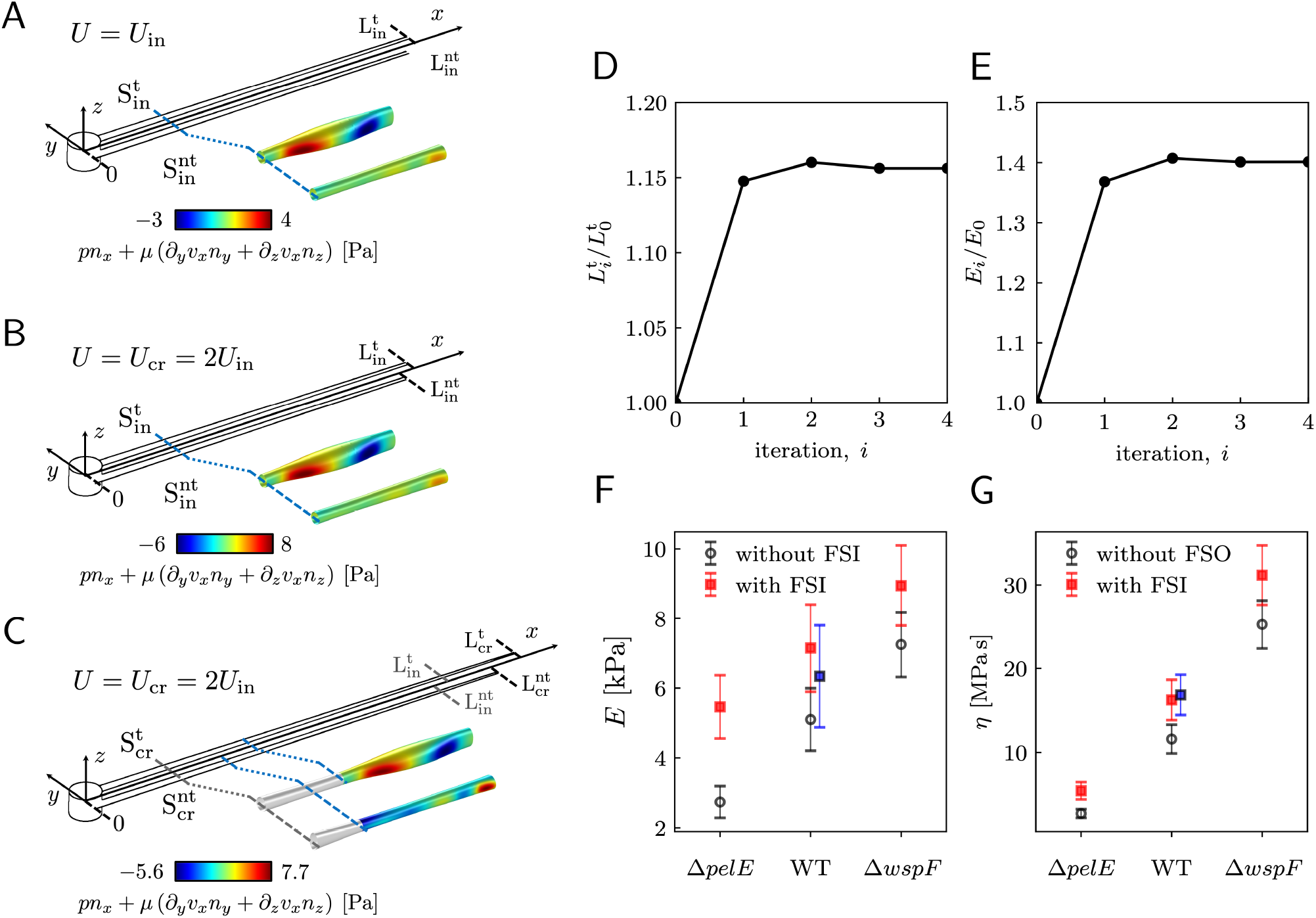
(A) Schematic of the streamers during the initial stage of the creep-recovery test. The zoomed-in portion of streamers shows *f*_in_ (*x*) (Eq. (7)) in the region between *x* = 810 μm and *x* = 870 μm. *f*_in_ (*x*) is calculated with CFD using the initial shapes 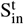 and 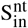 of the tracked and not tracked streamers, respectively, and *U*_in_. (B) Schematic of the streamers during the creep stage of the test, when FSI is neglected. CFD simulations with the initial shapes 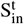 and 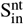 and *U*_cr_ = 2*U*_in_, gives *f*_cr_ (*x*) = 2 *f*_in_ (*x*), at low Re. (C) When taking FSI into account, *f*_cr_ (*x*_cr_ (*x*,*t*)) changes with time, because it depends on the time-dependent shape of the filaments 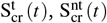 (Eq. (12)). Our estimate of *f*_cr_ (*x*_cr_ (*x*,*t*\*)) with FSI was obtained by calculating the approximate shapes 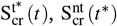 at time *t* = *t*^*^ ≡ 150 *s*, right after the elastic deformation. *f*_cr_ (*x*_cr_ (*x*,*t*^*^)) is not trivially related to *f*_i*n*_ (*x*). We also see that the deformation is not the same on the two filaments, due to their different morphology. (D) Convergence plot for the elastically deformed length *L* and (E) *E* of the average PA14 WT streamer. (F,G) *E* and *η* calculated without FSI (open circles) and with the FSI correction calculated on the average streamer shapes (red squares). Blue squares for PA14 WT are obtained by calculating E and *η* with FSI for each experimental replicate and then by averaging the results.

We used this result to calculate *E* and *η* of 15 h-old streamers formed by *P. aeruginosa* PA14 WT. The integrals in Eqs. (3) and (4) were numerically computed with custom Python software, by discretizing them onto an equally spaced grid of 1 μm. Results from 20 independent experimental replicates show that *P. aeruginosa* PA14 WT produces streamers with Young’s moduli of the order of *E* ∼ 10^3^ Pa and viscosities of the order of *η* ∼ 10^7^ Pa s. The mean values and standard deviations of the mean for *E* and *η* are reported in Table and plotted in Figs. 5F and 5G (WT, open circles).

We estimated the average relative contribution of the elastic and viscous deformations during the creep-recovery tests. The elastic deformation 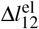 represents the main contribution, being on average 94% of the total deformation Δ*l*_12_ (*t* = 450 s) accumulated at the end of the creep stage.

### 3.5 Estimating the effects of fluid-structure interaction

In the previous paragraph, we calculated the axial stress *σ*_cr_ (*x*) on the streamers during the creep stage by assuming that the flow boundaries, namely the surfaces 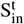 and 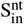 of the streamers, are not dependent on time. We then used this estimate of the axial stress to analyze the time dependent deformation of the streamers and to obtain the rheological parameters. With this approximation, we neglected the fact that during the creep stage the axial stress does not change only because *U* increases, but also because the filaments are stretched and the surface exposed to flow is deformed (Fig. 5C). In order to go beyond this approximation, we now present a method to estimate the effect of the moving boundaries on the hydrodynamic stresses and to correct the rheological results according to FSI.

During the creep stage, the time-dependent shape of a streamer can be written as:

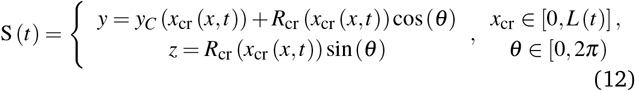

where *x*_cr_ (*x*, *t*) is the position at time *t* of the infinitesimal streamer element initially located at *x* and *R*_cr_ (*x*_cr_ (*x*,*t*)) is its deformed radius at time *t*. Thus, if we consider FSI, the time-dependent stress on the infinitesimal element of streamer initially located at *x* can be written as:

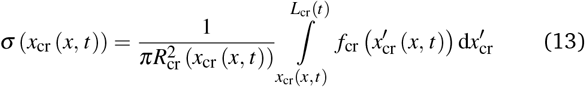

where d*x*_cr_ = (1 + *ε* (*x*,*t*)) d*x* is the deformed length of the infinitesimal element at time *t*. Even at low Re, the relation between *f*_in_ (*x*) and *f*_cr_ (*x*_cr_ (*x*,*t*)) is not a simple proportionality, since not only the average flow velocity doubles, but also the flow boundaries change with time when the two filaments are stretched. As reported at the end of the previous paragraph, the elastic deformation represents the main contribution to the flow-induced deformation. Thus, in order to estimate the effect of FSI, we can calculate an approximate value of the axial stress 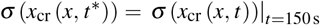 right after the elastic deformation at *t* = *t*\* ≡ 150 s. The shapes 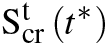 and 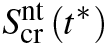 after the elastic deformation can be estimated by following an iterative scheme. The inputs for the first iteration are the initial shapes 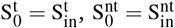 of the two filaments and the Young’s modulus 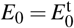 of the tracked filament, calculated without FSI, according to section 3.4. At the *i*-th iteration (*i* = 1, 2,..., *n*), we estimate the elastically deformed filament shapes 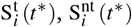 by calculating the strain *ε*_*i*_(*x*, *t*\*) of each filament as a function of *x*, according to

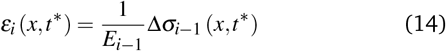

Here we assume that both the filaments are characterized by the same value 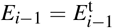 of the Young’s modulus, constant along their length, which is experimentally measured from the deformation curve of the tracked filament. In order to update the shape of the filaments at each iteration, we assume that, after the elastic deformation, the radius *R*_*i*_ (*x*) of each infinitesimal element changes according to the conservation of volume:

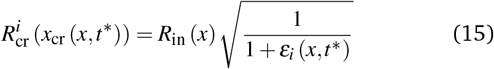

This corresponds to assuming a Poisson’s ratio *ν* = 0.5 for the streamers, which is within the range of values reported in literature for surface attached biofilms. ^41–44^ With the new morphologies 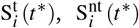, we perform a numerical simulation and obtain a new estimate of 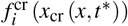. We then calculate the axial stress *σ* (*x*_cr_ (*x*, *t*\*)) for both filaments according to Eq. (13), where *x*_cr_ (*x*,*t*\*) is calculated as 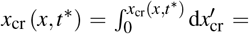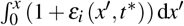. Then, by using *σ*_*i*_ (*x*_cr_ (*x*,*t*\*)), we calculate the axial stress step Δ*σ*_*i*_ (*x*, *t*\*) = *σ*_*i*_ (*x*_cr_ (*x*, *t*\*)) − *σ*_in_ (*x*, *t*\*). Finally, *E*_*i*_ and *η*_*i*_ are obtained according to Eqs. (3) and (4). We repeat this iteration scheme until the difference between subsequent values of *E*_*i*_ and length 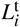 for the tracked filament is smaller than 1% (Figs. 5D and 5E). During the stress tests, we cannot measure the time-dependent deformation on the whole length of the filaments. So, in order to check for the consistency of the iteration results, we verified that the elastic deformation of the tracked portion of streamers is compatible with the one computed iteratively.

In order to estimate the average weight of this FSI correction, we applied the iterative scheme to a streamer model built with the average Young’s modulus (Table 1) and the shape averaged over all the 20 experimental replicates. To calculate the average shape, first we rescaled the initial shapes *S*_0_ obtained in each experimental replicate to the average length *L*, and obtained:

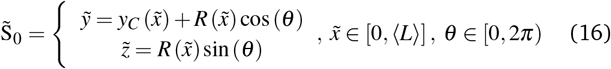

where 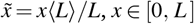. Then we averaged the resulting 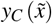 and 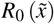 over the replicates. The convergence plots for the iteration scheme applied to this system are reported in Figs. 5D and 5E. The average length of the 20 replicates used for the rheological study was *L* = 1.99 mm. The final value for the length is *L*_*n*_ = 2.3 mm, a factor 1.16 higher than the initial length. The elastic strain of the whole filament is larger than the one observed on average for the portion of streamer tracked during the stress tests, reported in section 3.3. This is due to the fact that the non-homogeneous strain *ε*_el_ (*x*) (Eq. (2)) has typically its maximum downstream of ROI_2_. The rheological parameters after the FSI correction are *E* = 7.2 ± 1.3 kPa and *η* = 16.3 ± 2.4 MPas (Figs. 5F and 5G, red squares). Thus, neglecting FSI underestimates the applied stress, and thus the elasticity and viscosity, of a factor 1.4.

**Table 1.** Mean values and standard deviations of the mean for *E* and *η* calculated over 20 independent experimental replicates of 15 h-old streamers of *P. aeruginosa* PA14 WT.

| | average | $\sigma_{mean}$ |
| --- | --- | --- |
| $E$ [kPa] | 5.1 | 0.9 |
| $\eta$ [MPa s] | 11.6 | 1.7 |

We compared the correction estimate on the averaged streamer with the values obtained by applying the iteration scheme to each one of the 20 experimental replicates and then by averaging the corrected results. In this case, we obtained *E* = 6.3 ± 1.5 kPa and *η* = 16.9 ± 2.4 MPas (Figs. 5F and 5G, blue squares). The results obtained with the two averaging procedures are compatible within the error bars. However, performing the correction procedure on the average streamers is a computationally cheaper option, since the time required to perform the iterations scales as the number of replicates. To conclude this section, we have to point out that some of the samples we tested showed small discrepancies between the initial elastic elongation and the deformation recovered at the end of the creep stage. We verified that such discrepancies are compatible within the correction for FSI.

### 3.6 Rheological characterization of streamers with different composition

Our technique can detect differences in the rheology due to differences in biochemical composition. We compared streamers grown by PA14 mutants differing in the production of Pel, the primary polysaccharide in the matrix of PA14:^45^ PA14 WT, the Pel deficient mutant PA14 Δ*pelE* and the Pel overproducer strain PA14 Δ*wspF*. As we discussed in a recent article, ^16^ the selective staining of eDNA and Pel allows the characterization of the biochemical composition of the streamers and of the distribution of such components throughout the filaments. The differences in the biochemical composition of the streamers are reflected in both their morphology (〈*L*_WT_〉 = 2.22 ± 0.08 mm, 〈*L*_ΔpelE_〉 = 2.87 ± 0.13 mm and 〈*L*_ΔwspF_〉 = 1.36 ± 0.06 mm) and rheology (Figs. 5F and 5G, open circles). The typical experiment for comparing the three strains was performed by simultaneously flowing the bacterial suspensions in different channels of the same device, in order to minimize biological variation. We repeated this experimental protocol five times, with independent bacterial batches. We used the average morphological and rheological results (ESI†, Table S1) as the initial values for the iterative scheme to quantify the impact of FSI. The resulting FSI correction is equal to a factor 2 in the case of PA14 Δ*pelE*, 1.4 in the case of PA14 WT, and 1.2 in the case of PA14 Δ*wspF* (Figs. 5F and 5G; ESI†, Table S1). The difference in the FSI correction factor is attributable both to the difference in morphology and rheology: on the one hand, for given *E* and *η*, longer streamers will undergo a higher deformation; on the other hand, for a given length, the stiffer the filaments, the smaller the deformation and, consequently, the relevance of the FSI correction. In conclusion, our iterative scheme makes it possible to easily take into account the FSI impact on the measured values and to reliably compare streamers with different morphologies and rheology, formed by different bacteria and in different growth conditions.

## 4 Discussion and conclusion

In this paper, we present a microfluidic platform that allows the formation of biofilm streamers in a highly reproducible way and the systematic characterization of their morphology, biochemical composition and rheology *in situ*. Previous works using micropillar arrays focused on the formation of intricate webs of streamers,^12,14,15,46–48^ while isolated thin filaments were observed around bubbles ^38^ or oil droplets. ^27^ A fine control of the hydrodynamic conditions around the isolated PDMS pillars in our device lead to the nucleation of pairs of parallel, straight streamers. The selective fluorescent staining of the EPS ^13^ can be exploited so as to characterize not only the biochemical composition of the streamers, but also their morphology. We verified that this approach provides a finer resolution of the morphological details than that achieved by observing only the few cells attached to the streamers.^12,15,17,27,29,40^ In our configuration, the streamers are typically much more extended than the region covered in bacterial cells. Moreover, fluorescent staining of the EPS allows the visualization of the streamers without altering their structural properties, as embedding tracers would do in thin filaments.^14,38,46–48^ In this study, we exploited eDNA staining in order to visualize the streamers and to obtain a detailed characterization of their morphology. This visualization approach is crucial for the reliability of the hydrodynamic and rheological characterization of the system, since the flow field and the force exerted on the streamers depend to a great extent on the details of their morphology. The detailed morphological information allowed us to perform accurate 3D CFD simulations in order to determine the fluid dynamic conditions in each experimental replicate. Thus, the possibility of taking into account the actual morphology of the streamers, and not just estimates of their average cross-sectional area, ^21–23,25^ allowed us to improve the quantification of the mechanical properties of a portion of streamer tracked during a creep-recovery test. Additionally, the PTV experiments confirmed the accuracy of the no-slip boundary conditions and the circular approximation for the cross section of the filaments used in the simulations. Thanks to the precise characterization of the streamer morphology and flow field, we can exploit our platform to estimate the drag increase caused by the streamers directly. By comparing the drag force on the pillar before and after streamer formation, we quantify an increase of about an order of magnitude. In the context of marine particle dynamics, previous studies ^17,18,27^ estimated the drag increase after streamer formation by considering only a limited portion of the streamers in the vicinity of the particle. Thus, the calculations performed by neglecting the whole length of the streamers underestimate the actual drag increase. Our platform makes it possible to overcome such limitations and obtain new insight on the transport of colonized particles, such as marine snow or oil droplets.

In a recent research study performed using this microfluidic platform, ^16^ we were able to precisely characterize *Pseudomonas aeruginosa* streamers, and find a mechanistic link between their biochemical composition and their structural and rheological properties for the first time. This uncovered the role of eDNA as the fundamental building block of *P. aeruginosa* streamers, a finding with important implications in groundwater and marine research and filtration systems. The precise control of microfluidics over microenvironmental conditions will also make it possible to compare the effects of different physico-chemical conditions, such as pH, temperature or ambient flow on streamers development. The role of ambient flow in determining the properties of biofilm streamers is particularly interesting, since streamer formation and integrity are intimately linked to the hydrodynamic conditions in the surroundings. Interestingly, flow-driven aggregation has been observed even in abiotic systems, where a suspension of particles and polymers formed abiotic streamers while flowing through an array of micropillars. ^49^ This suggests that our platform could potentially find applications in the study of flow-driven aggregation in a variety of soft matter systems.

## Author Contributions

E.S., R.R. and R.S. designed research. G.S. performed the experiments, analyzed all the data, designed and built the mathematical model. J.S., E.S. and R.R. contributed to the design of the mathematical model. G.S. and E.S. wrote the manuscript and all authors edited and commented on the manuscript.

## Conflicts of interest

There are no conflicts to declare.

## Acknowledgements

The authors acknowledge Dr. Sam Charlton, Steffen Geisel, Prof. Luca Heltai, Dr. Alberto Sartori, and Prof. Jan Vermant for the insightful discussions; Ela Burmeister for the technical support; Prof. Leo Eberl for providing the P. aeruginosa PA14 mutant strains; support from SNSF PRIMA grant 179834 (to E.S.), from SNSF Ambizione grant PZ00P2_202188 (to J.S.), from Gordon and Betty Moore Foundation Marine Microbial Initiative Investigator Award GBMF3783 (to R.S.), and from Simons Foundation Grant 542395 (to R.S.) as part of the Principles of Microbial Ecosystems Collaborative (PriME).

## Electronic Supplementary Information (ESI)

### 1 Axial stress on a streamer in a background flow

According to ^1^, the stress tensor *σ*_*ij*_ in an isotropic, Newtonian fluid can be written as:

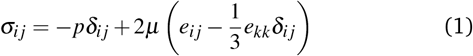

where *p* is the pressure, *μ* is the dynamic viscosity of the fluid and *e*_*ij*_ is the rate of strain tensor, defined as

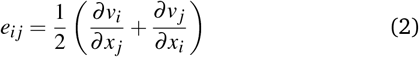

We can consider our liquid medium as incompressible, so *e*_*kk*_ = 0 and Eq. 1 takes the form:

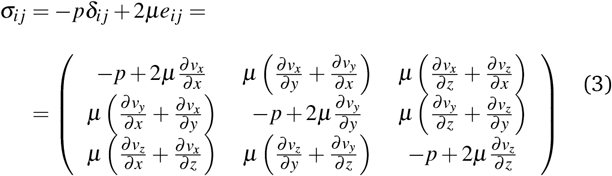

Thus, the force per unit area on the infinitesimal surface with normal 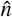 is:

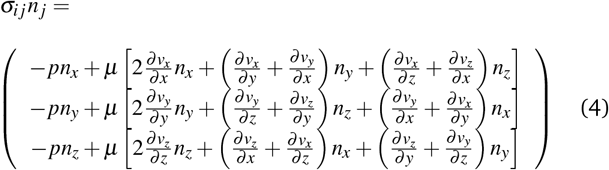

The *x* component is:

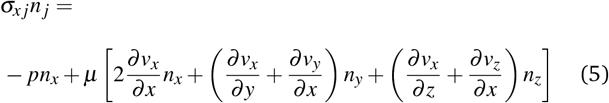

Given the typical shape of the streamers and the flow field in our device we have that in the region from *x* = 400 μm to the end of the streamer 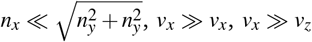 and the velocity field varies slowly with x so that *∂*_*x*_ ≪ *∂*_*y*_ and *∂*_*x*_ ≪ *∂*_*z*_. Under such conditions, we can safely write:

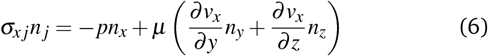

Thus, the force in the x-direction exerted on the infinitesimal surface d*A*_S_ with normal 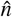 is *σ*_*ij*_n_*j*_ d*A*_S_. The general expression for d*A*_S_ for a streamer with variable circular cross section with radius *R* (*x*) is 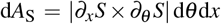, where *S* = *S* (*x*, *θ*) is the parametrization of the surface of the streamer given in Eq. 1. The magnitude of the cross product can be calculated by considering *S*: *R*^2^ → *R*^3^ as:

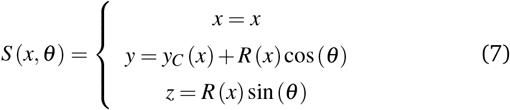

Its derivatives are

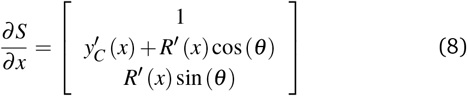

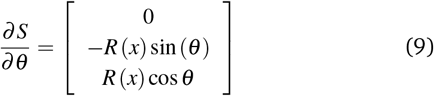

Thus, the cross product is

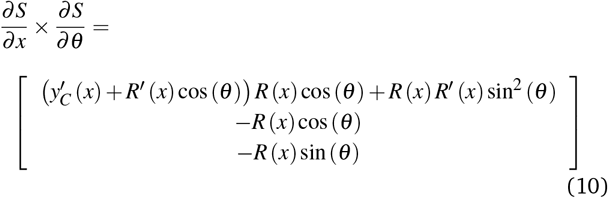

By taking the magnitude of the resulting vector, we obtain:

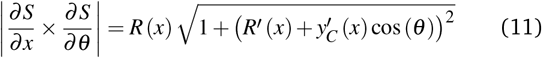

and for the area of the infinitesimal surface element:

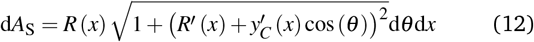

With the typical shape of the streamers in the region from *x* = 400 *μ*m to the end of the streamer, the *R* (*x*) and *y*_C_ (*x*) vary slowly with *x*. We can thus approximate Eq. 12 as 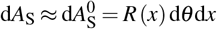.

**Fig. S1.**
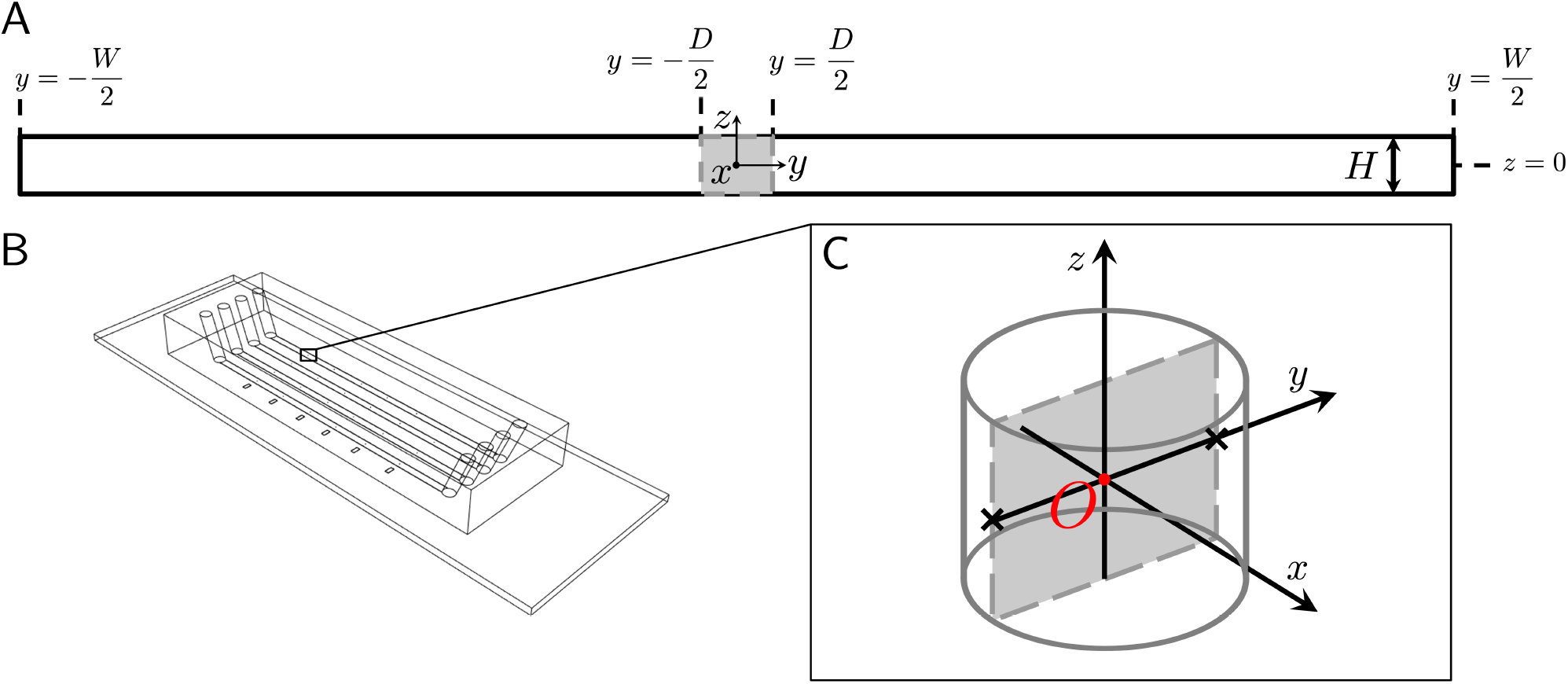
(A) Cross-sectional view of a channel, cutting a in half a pillar (shaded region). The flow direction is aligned with the *x* axis, pointing out of the image plane. The origin of the frame of reference is at the center of the pillar. The channel walls are located at *y* = −*W*/2 and *y* = *W*/2. The height of the channel is *H* = 40 μm. (B) Schematic of the microfluidic platform, containing four channels, with six pillars inside each. The position of the pillars are marked by six notches patterned at the side of the device. The streamwise inter-pillar distance is 5 mm. The channels have independent inlets and outlets. (C) The coordinate system used to describe the pair of streamers tethered to a pillar in our device. The origin *O* (red point) is located at center-width and half-height of the channel, at the center of mass of the cylinder. The *x* axis is aligned with the direction of flow. The tethering points of the two streamers lie on the *y* axis, at *y* = −*D*/2 and *y* = *D*/2 (black crosses).

### 2 Meshing sequence

CFD simulations were performed by using a mesh with local refinements in the regions near the pillar and the streamers (Fig. 2A). To this end, we built subdomains around them. The length of each subdomain in the *x* direction was 80 μm. We obtained fine and high-quality elements around the streamers with a swept mesh. For the source face we used a free triangular mesh (Fig. 2B, color map, colored according to element quality), which was swept along the length of the filaments to mesh the other subdomains (Fig. 2B, gray subdomains). In each subdomain, the mesh was swept on 80 layers, so that each element had a 1-μm side in the *x* direction. The subdomains with the pillar and the end of the filaments, which are not meshable with a sweep operation, were meshed with an extra fine free tetrahedral mesh. The rest of the channel were meshed with a free tetrahedral mesh with coarse elements. Element quality was calculated by using a COMSOL built-in estimator based on element skewness as:

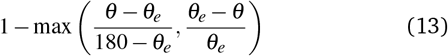

Where *θ* is the angle over an edge in the element, *θ*_*e*_ is the angle of the corresponding edge in an ideal element, and the maximum is taken over all edges of the element. The quality of a perfectly regular element is 1, while the quality of a degenerate element is 0.

**Fig. S2.**
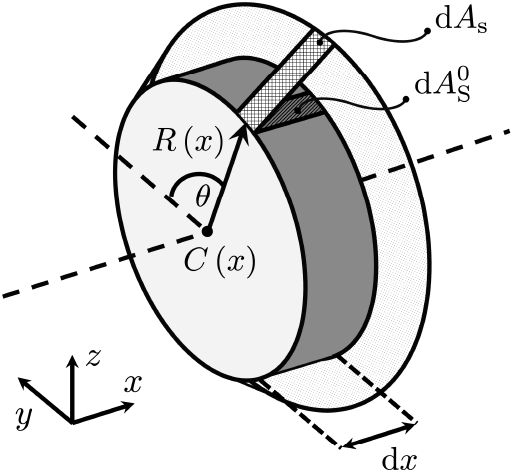
Approximate and true areas of an arbitrary infinitesimal surface element. The gray disk, located at *x*, with thickness d*x* and radius *R*(*x*), corresponds to the one shown in Fig. 2A. The area 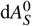 of the infinitesimal surface element of such a disk is 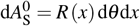. The true infinitesimal element of streamer would not necessarily be cylindrical and its area d*A*_*S*_ would be 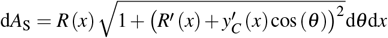

**Fig. S3.**
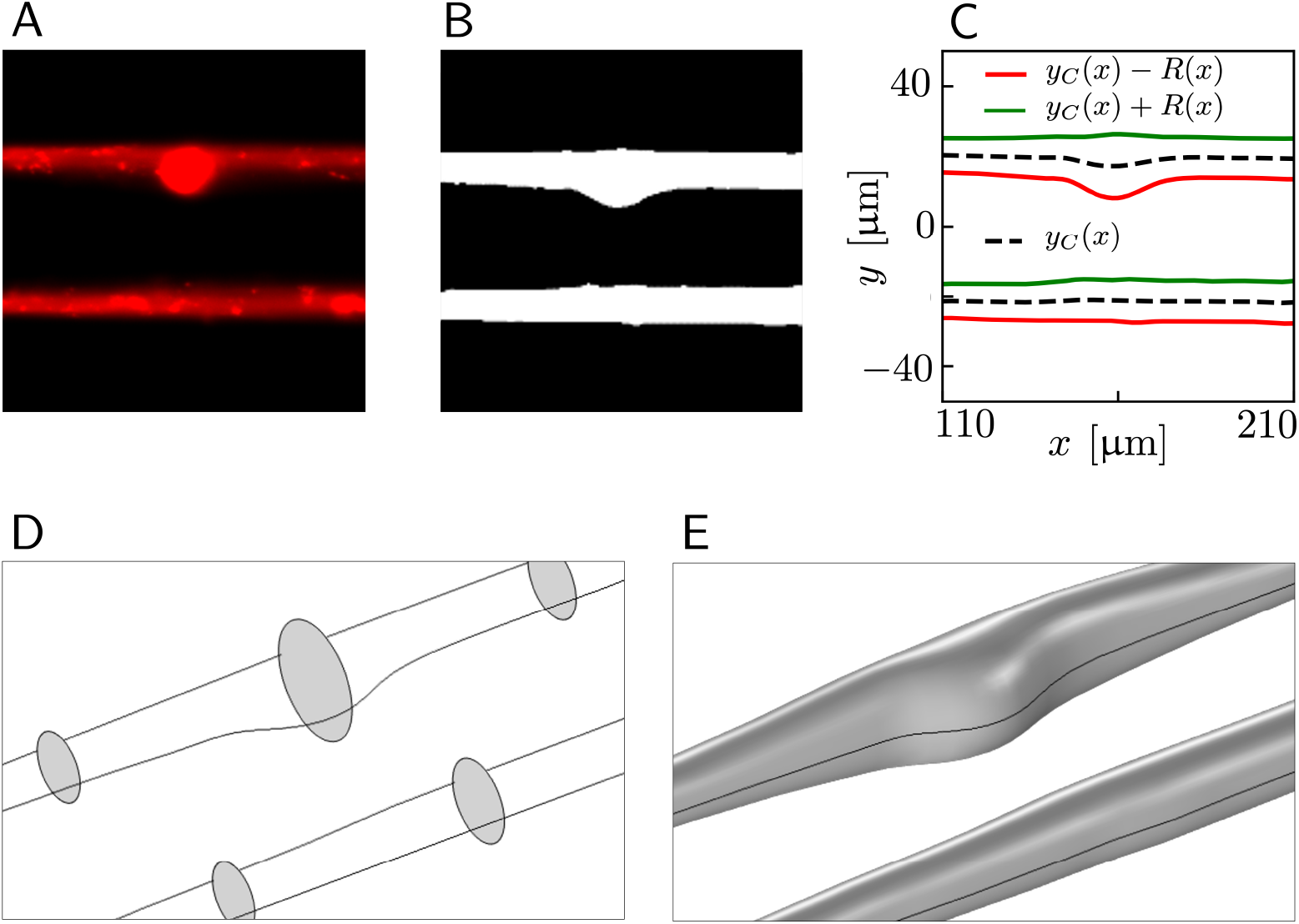
Workflow for building the 3D model for CFD simulations. (A) Fluorescence image of the streamers. (B) Detail of the binarized image with threshold 15%. (C) Detail of the shape of the streamers after interpolation and smoothing of the binarized data. (D) Circular cross sections and streamers outlines drawn on the channel midplane. The outlines are used as guide curves for interpolating the cross sections all along the length of the streamers. (E) Detail of the 3D geometry resulting from the loft operation.

**Table 1.**
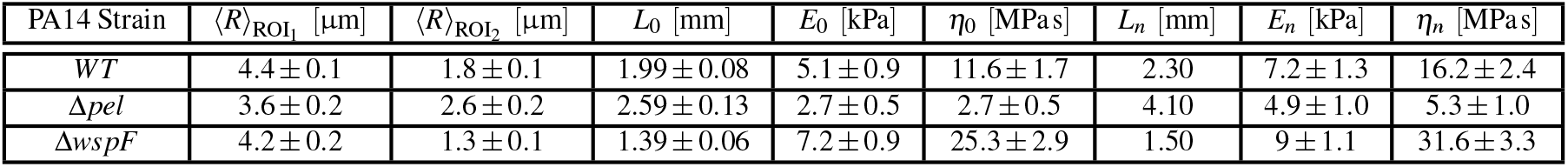
Morphological and rheological properties of the streamers formed by the three strains of PA14 compared in this study. The mean values and standard deviations of the mean were calculated by averaging the results obtained with five experimental replicates, performed with independent bacterial batches. In each experimental replicate we simultaneously flowed the cells of the three strains in different channels of the same device, in order to minimize biological variation. Altogether, we tested 20 independent streamers for each strain. *L*_0_, *E*_0_ and *η*_0_ are the values obtained without FSI. *L*_*n*_, *E*_*n*_ and *η*_*n*_ are the values that take FSI into account, obtained at the end of the iteration procedure (at the *n*-th iteration step). The convergence was reached after *n* = 4 iterations in the case of PA_14_ WT, after *n* = 7 in the case of PA14 Δ_*pel*_, and after *n* = 3 in the case of PA14 Δ_*wspF*_

**Fig. S4.**
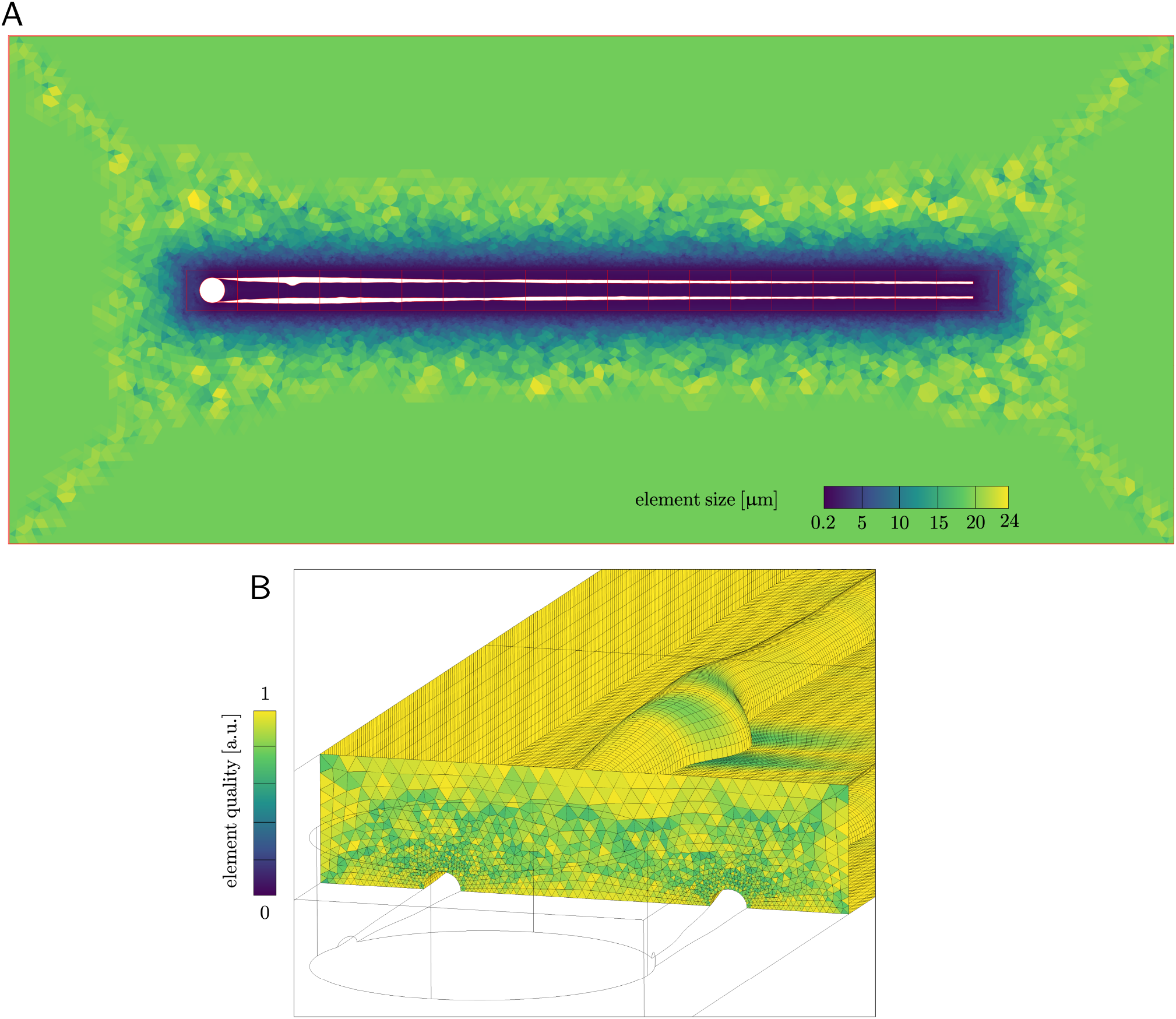
(A) Element size map on the midplane. (B) Map of element quality on the source surface for the loft operation and on the surfaces of the swept domains

**Fig. S5.**
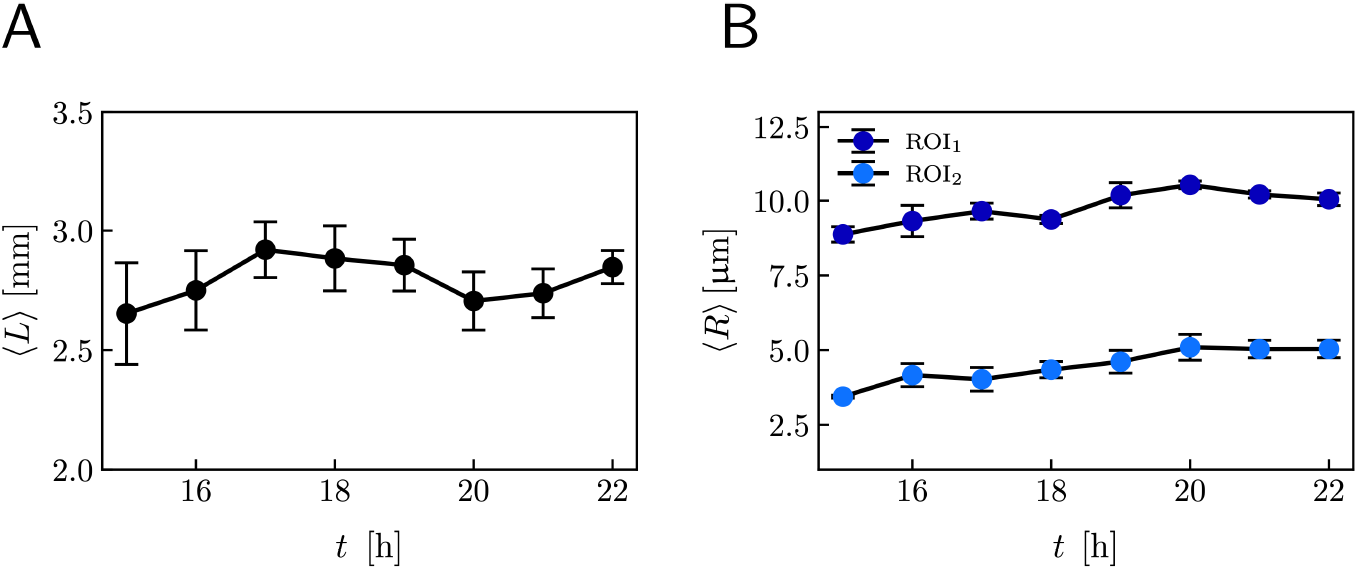
Representative data of length and radii on ROI_1_ and ROI_2_ (Fig. 1A) as a function of time between *t* = 15 *h* and *t* = 22 *h*. We observe that within this time interval, the morphology of the streamers typically does not change significantly. The reported data were averaged over 4 streamers.

**Fig. S6.**
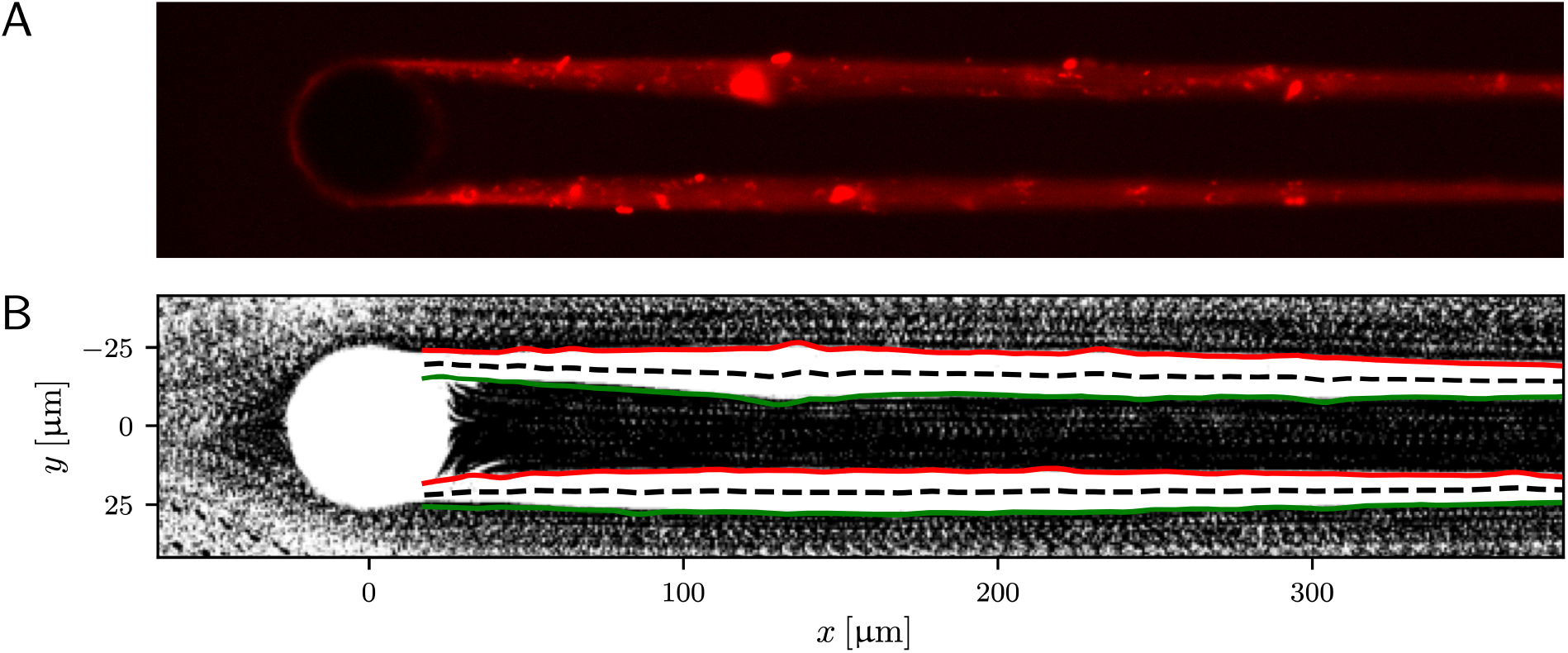
(A) Fluorescence image of PA14 wt streamers. (B) Outline of the streamers shown in panel A, obtained as explained in section 2.3, superimposed to the minimum intensity projection of a stack of images of tracers acquired for PTV. PTV images were processed as explained in section 2.4.2 to select only particles flowing on the channel midplane. The regions of the pillar and the filaments are white because no particle flows through throughout the whole acquisition. This proves that the fluorescence signal provides a good characterization of the streamers’ morphology.

## Notes

### Competing Interest Statement

The authors have declared no competing interest.

